# *IIIandMe*: An Algorithm for Chromosome-scale Haplotype Determination Using Genome-wide Variants of Three Haploid Reproductive Cells

**DOI:** 10.1101/2022.12.07.519546

**Authors:** Han Qu, Ruidong Li, Lei Yu, Weiming Chen, Yuanfa Feng, Qiong Jia, Ryan Traband, Xuesong Wang, Shibo Wang, Luoxian He, Zixian Wang, Meng Qu, Sergio Pietro Ferrante, Jianguo Zhu, Weide Zhong, Mikeal Roose, Zhenyu Jia

## Abstract

Our recent algorithm, *Hapi*, infers chromosome-scale haplotypes using genomic data of a small number of single gametes. Its advanced version, *IIIandMe*, is proposed here to achieve comparable phasing accuracy with as few as three gametes, pushing the analysis to its limit. The new method is validated with simulation and a citrus gamete dataset. The rapid advances in genotyping technologies promise a broad application of *IIIandMe* in disclosing important genetic information.

## Main Text

Mounting evidence has shown the benefit of using haplotype variants over single nucleotide polymorphisms (SNPs) in various genetic analyses, including genome-wide association studies (Yang et al. 2012; Howard et al. 2017; Lambert et al. 2013; Zhang et al. 2021), detection of the signatures of positive selection (Fariello et al. 2013), deducing genetic admixture, introgression, and demographic history (Lohmueller, Bustamante, and Clark 2009; Palamara et al. 2012). Accurate chromosomal haplotypes are needed to identify causal haplotype variants for further genetic dissection. We recently developed the *Hapi* algorithm to infer chromosome-length haplotypes using the genotypic data of several single gametes (R. Li et al. 2020). The genomic data of gametes, represented by variants at heterozygous SNP (hetSNP) loci along the individual genome, may be obtained by various platforms, including whole genome sequencing or genotyping arrays. Multi-stage preprocessing steps, including removal of erroneously genotyped markers and iterative imputation of missing markers, are implemented by *Hapi* when the quality of gamete data is suboptimal. With the rapid advancement of biotechnologies, high-resolution genotyping with negligible errors will be achieved in the foreseeable future. Here we present a new and advanced chromosome-phasing algorithm, *IIIandMe*, which only requires three gametes when genotypic data are of high quality. Compared to *Hapi*, *IIIandMe* is logically more straightforward and computationally more efficient.

We illustrate how *IIIandMe* works with three haploid gametes given the diploid genotype of hetSNPs of the donor, where two different colors (green and yellow in **Figure 1**) represent two complementary (or reciprocal) parental haplotypes for a chromosome. The same principles apply to phasing other chromosomes. A few prerequisite rules (or assumptions) are needed for the subsequent reasoning and computation described in *IIIandMe*. Rule {1} – DNA breakpoints due to crossovers are randomly positioned (between two adjacent hetSNPs) along any haploid gamete chromosome. Rule {2} – Theoretically, there exists a complementary haploid chromosome of any observed haploid chromosome (recombinant or nonrecombinant) in a given gamete; these two reciprocal chromosomes may be considered as the product from the same meiotic event. However, it is very unlikely (probability close to zero) to sample these two reciprocal gamete chromosomes since the sample space (the population of all gametes from the donor) is hypothetically large. Rule {3} – Based on rule {2}, the probability that two sampled gamete chromosomes having breakpoints at the same position is close to zero. This is likely to be true if marker density is substantially higher relative to the frequency of crossovers. Rule {4} – If two gamete chromosomes in a sample have complementary or identical genotypes, then both must represent the intact parental haplotypes (without breakpoint). Note that rule {4} is a necessary consequence of rules {1} to {3}.

**Figure 1.**
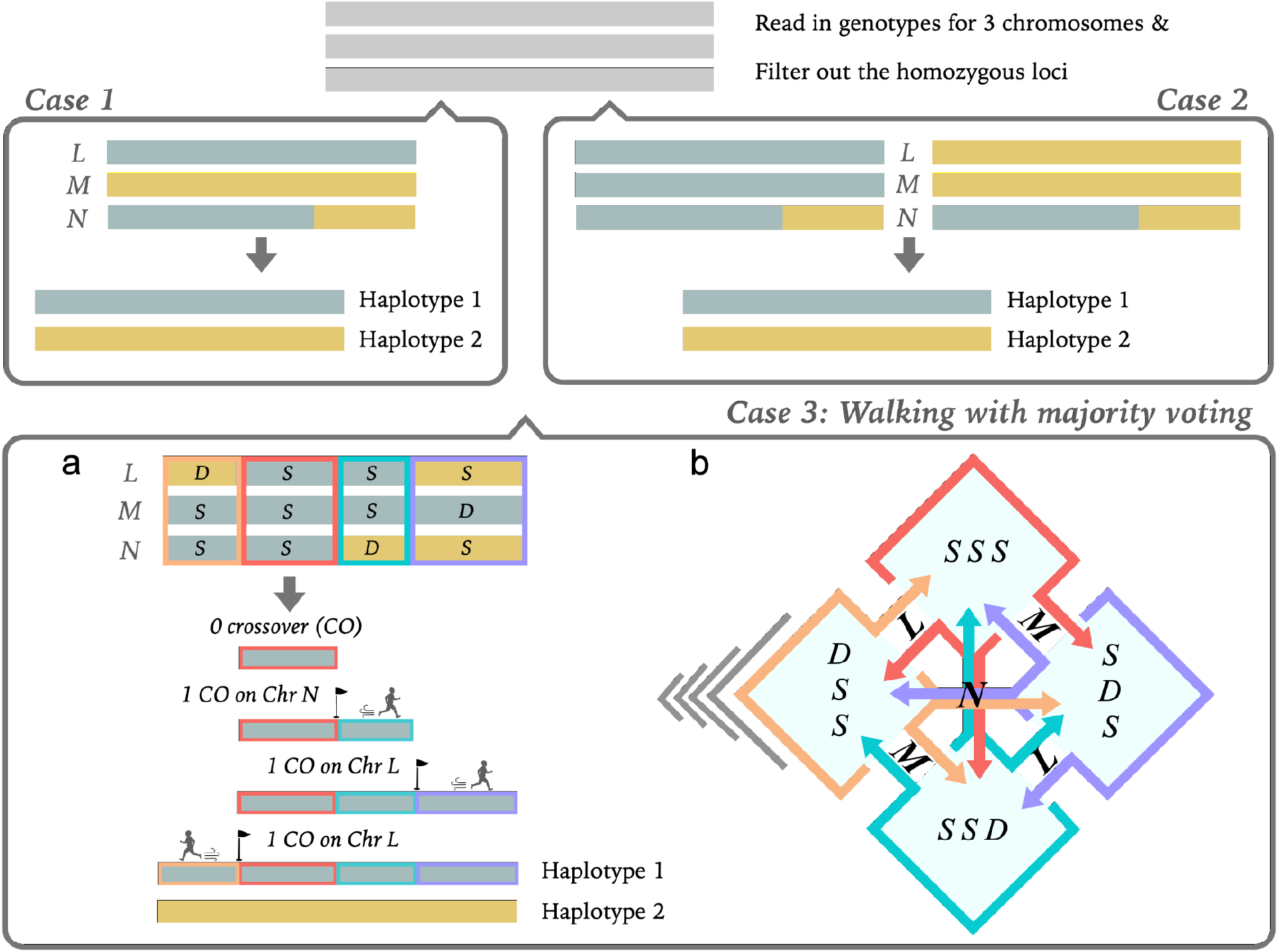
The rationale of the core 3-gamete algorithm implemented by *IIIandMe*. **Cases 1 & 2:** The genotypes of two out of three gamete chromosomes are complementary or identical, respectively. Therefore, the parental haplotypes can be immediately obtained. **Case 3:** Two or three gamete chromosomes have breakpoints. **(a)** The workflow of implementing one-locus-a-step walking with majority voting (*WMV*). A common region (*S*-*S*-*S*) is first identified, followed by the consequent detections of crossovers (labeled with a flag) on both sides of this common region. Haplotype 1 is first inferred and haplotype 2 can be obtained by flipping haplotype 1. **(b)** Pattern-transition diagram facilitating the detection of the gamete chromosome with a crossover. The red, orange, blue and purple boxes represent *S*-*S*-*S*, *D*-*S*-*S*, *S*-*S*-*D* and *S*-*D*-*S*, respectively. The color arrows denote the pattern-to-pattern transition at a locus with the detected crossover and the associated letters (*L*, *M* and *N*) indicate the chromosome with that crossover.

We start from the haploid genotypes for three gamete chromosomes, labeled with *L*, *M* and *N*, respectively (**Figure 1**). In the simplest scenarios with two nonrecombinant gamete chromosomes (shown in Cases 1 or 2), the parental haplotypes for that chromosome can be immediately obtained according to rule {4}. Panel **a** of case 3 represents a common scenario where at least two gamete chromosomes have breakpoints. A *walking with majority voting (WMV)* strategy is implemented, described as follows, to infer the parental phases of the chromosome. A common region, denoted by the pattern of *S*-*S*-*S* (red box), will be first identified across the three gamete chromosomes. If no such a common region can be found, we simply flip the entire genotype of one or two original gamete chromosomes to yield a common region (Online Methods). The word ‘flip’ refers to the swap between the two reciprocal genotypes at heterozygous loci (rule {2}). This common region, representing a phased fragment of one parental haplotype (Haplotype 1 in **Figure 1**), is then used as the backbone of Haplotype 1 from which we can extend on both sides through *WMV*. There are four genotype patterns for *L, M* and *N*, *i.e*., *S-S-S* (red box), *D*-*S*-*S* (orange box), *S*-*S*-*D* (blue box) and *S*-*D*-*S* (purple box), at each hetSNP locus, where *S* and *D* denote ‘same’ and ‘different’ genotypes, respectively. In the one-locus-a-step ‘walk’, a transition between any two genotype patterns indicates a crossover on only one of these three chromosomes based on the majority voting principle - the majority of gametic chromosomes do not bear crossovers at any locus. For example, a transition from *S*-*S*-*S* to *D*-*S*-*S* implies a crossover on chromosome *L*. In *WMV*, we monitor transfers between these genotype patterns to detect crossover-bearing chromosomes and infer the genotypes along Haplotype 1 using a pattern-transition diagram (panel **b** of Case 3). Haplotype 2, which is complementary to haplotype 1, can be simply obtained by flipping the genotypes of the inferred Haplotype 1.

If a rare scenario that two gamete chromosomes have a crossover at the same locus happens in this 3-gamete analysis, *IIIandMe* claims a crossover on the third nonrecombinant chromosome incorrectly because of the aforementioned majority-voting scheme. To preclude this potential error due to this very-low-probability event, we recommend slightly increasing the sample size, for example, to a set of 5 or 6 gametes. When analyzing 5 gamete chromosomes, there will be 10 (choose 3 out of 5) possible combinations of three gamete chromosomes. One can apply *IIIandMe* to each of these 10 combinations, yielding 10 sets of inferred haplotypes for the chromosome. The majority-voting will be conducted for one more time on these 10 sets of candidate haplotypes, yielding the consensus haplotypes for the chromosome. The *IIIandMe* program, which is publicly available at https://github.com/Jialab-UCR/IIIandme, will automatically implement a 3-gamete analysis or such a *k*-gamete analysis if *k*, the number of inputted gametes, is greater than 3.

A simulation using a maize dataset (Li et al. 2015) was performed to compare the performance of *IIIandMe* versus *Hapi* (online Methods). Compared to *Hapi, IIIandMe* had significantly reduced time consumption and RAM usage in the simulation (**Figures 2 A** and **B**). In the 3-gamete and 4-gamete analyses, neither method was able to correctly derive the parental haplotypes of chromosome 1, owing to the fact that two microspores have a common crossover (**Figure 2 C-3** and **C-4**). Nevertheless, in the 5-gamete and 6-gamete analyses, greater than 50% of the microspores do not have crossover at this locus so the two rounds of majority voting were able to identify these two chromosomes with a common crossover and inferred the parental haplotypes of chromosome 1 accurately (**Figure 2 C-5** and **C-6**).

**Figure 2.**
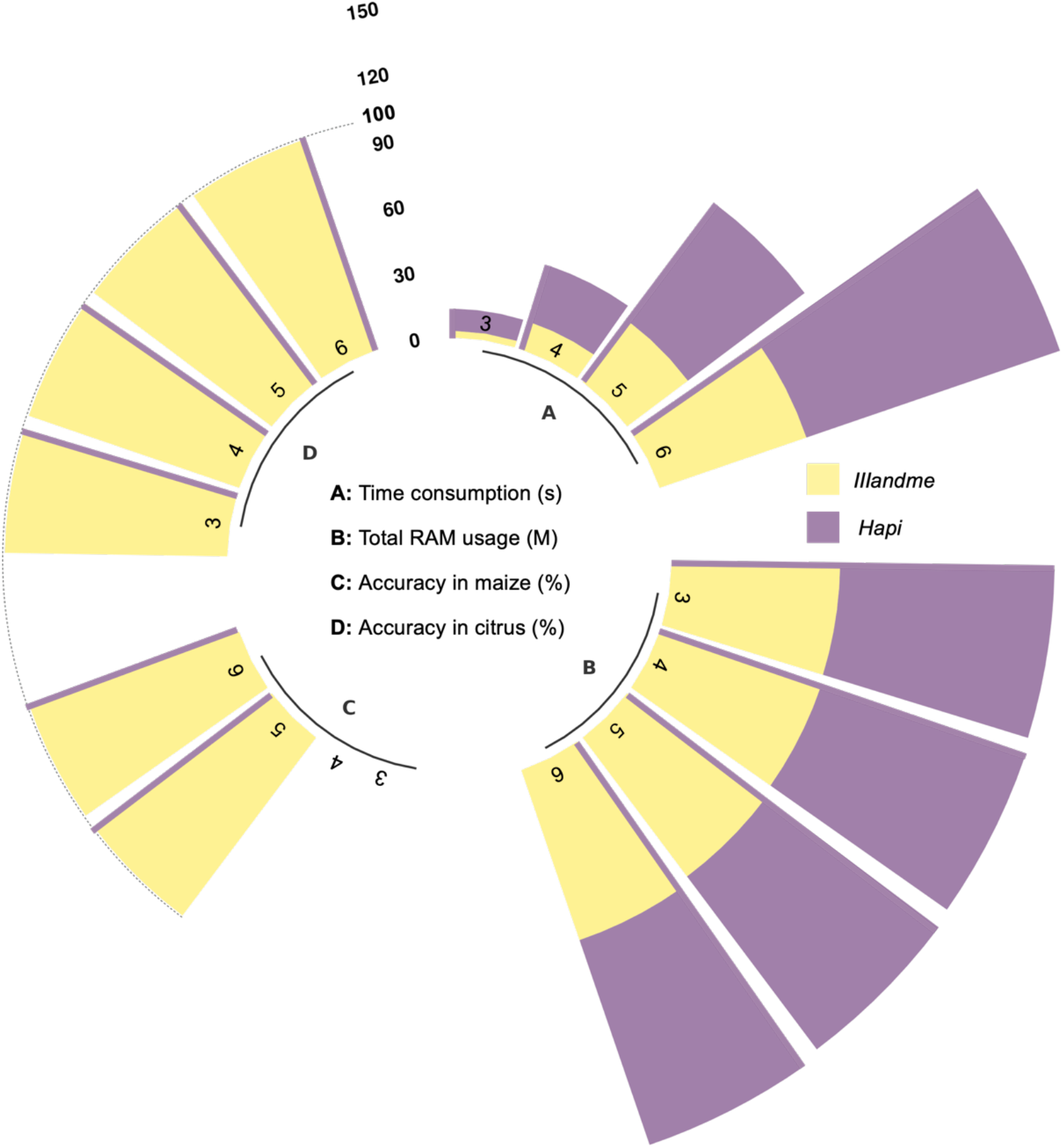
Performance comparison between *IIIandMe* and *Hapi* when analyzing 3, 4, 5 or 6 single gamete cells, respectively, in the simulation (**A**, **B** and **C**) and in the analysis of a citrus dataset **(D)**.

Our citrus dataset of *Clementine de Nules*, consisting of 6 single pollen grains, was analyzed for evaluating *IIIandMe* and *Hapi*. When analyzing any set of 3 pollen grains, 20 candidate haplotypes (choose 3 out of 6) were estimated for each of 9 chromosomes by each method. As shown in **Figure 2 D-3**, both methods were able to infer the parental haplotypes with 100% accuracy because no two or more pollen grains carry a common crossover at any loci. The inference accuracies for 4-gamete, 5-gamete, and 6-gamete analyses were also 100%, as displayed in **Figure 2 D**.

We demonstrated that *IIIandMe*, which applies to genomic data of high quality, can accurately infer chromosomal haplotypes using three or a few more single gametes and is computationally much more efficient than *Hapi*. Given that crossovers are very rare (Beye et al. 2006; Lyu et al. 2022), three gametes are theoretically sufficient for phasing the entire genome, likely pushing the boundary to its possible limit. We foresee that *IIIandMe* will find its way to become impactful in many genetic research areas and applications.

