## Supplemental information for "*IIIandMe*: An Algorithm for Chromosome-scale Haplotype Determination Using Genome-wide Variants of Three Haploid Reproductive Cells"

### Online Methods and Discussion

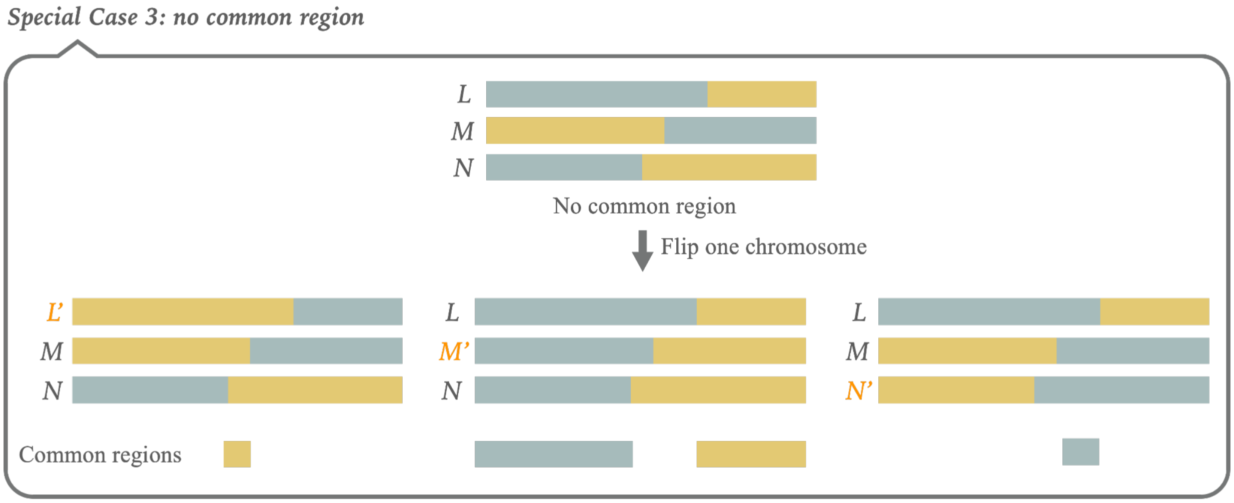


**Simulation:** Whole-genome sequencing data of 24 meiotic tetrads from a maize accession were initially published in (Li et al. 2015) and 24 'mutually independent' microspores were selected for developing *Hapi* (Li et al. 2020). A survey on the data of these 24 microspores showed that, on average, 1.015 microspores have a crossover at the same locus (between the same two adjacent hetSNPs), which is consistent with rule {3}. In the simulation, we selected chromosome 1 (82710 hetSNPs) of six microspores, in which two microspores have a crossover at the same locus. In each simulation scenario, we always included these two special microspores. Specifically, in a 3-gamete analysis with these two microspores, we randomly picked the third one from the rest of the four microspores, yielding four possible combinations (choose 1 out of 4) of a set of 3 microspores. The data of each combination were analyzed by *Hapi* and *IIlandMe*, respectively; the average performances for the two chromosome-phasing methods, including time consumption, total RAM usage, and inference accuracy, are presented in **Figures 2 A-3**, **B-3** and **C-3**, accordingly. In a similar way, the 4-gamete, 5-gamete, and 6-gamete analyses were performed with 6 (choose 2 out of 4), 4 (choose 3 out of 4), and 1 (choose 4 out of 4) combinations, respectively, and the average performances of respective methods are also summarized in **Figure 2**.

**Real data analysis:** We also demonstrated *IIlandMe* and *Hapi* using our dataset of a citrus accession, Clementine de Nules, including the diploid genotypic data for 540-2365 ordered hetSNPs **(Table S1)** on 9 chromosomes and haploid genotypic data for 6 single pollen grains. The cultivar has been assayed using a customized SNP array (AxiomTM Citrus56AX) that was designed at UCR by co-author Dr. Roose. This dataset has been deposited and is publicly available in the Citrus Genome Database (https://www.citrusgenomedb.org/).

| Chromosomes | The number of all SNPs | The number of hetSNPs |
|---|---|---|
| 1 | 6289 | 1019 |
| 2 | 7773 | 1479 |
| 3 | 10572 | 2365 |
| 4 | 6259 | 1319 |
| 5 | 6665 | 1665 |
| 6 | 5446 | 1348 |
| 7 | 4961 | 540 |
| 8 | 4608 | 884 |
| 9 | 5360 | 1314 |

Table S1. The number of all SNPs and hetSNPs of the citrus accession, *Clementine de Nules*

**Discussion:** A chromosome-level haplotype assembly pipeline, consisting of two algorithms *sgcocaller* and *comapr*, was recently developed for analyzing large-scale single gamete sequencing data (Lyu et al. 2022). This method was compared to *Hapi* and showed improved performance, likely due to their fine-tuned parameters and probability models in multiple steps which may be biased towards a large sample analysis. Including a large number of gametes tends to increase the resolution and completeness of the phasing results. Nevertheless, the accuracy of inferred haplotypes and recombination maps is not compromised when a small gamete sample is used (shown in our data), owing to the fact that the average number of crossovers on a gametic chromosome is less than 3 across species (Beye et al. 2006). We were not able to include this haplotype assembly method in our comparison because this method only uses raw cell-barcoded sequencing data but is not applicable to SNP genomic data from other platforms, including genotyping arrays used in our study.

Our goal is to reduce the sample size substantially, and therefore operational cost, for phasing individual genomes, such that population-based genetics and clinical genetics studies become affordable and feasible. We demonstrated that *IIIandMe*, which applies to genomic data of high quality, can accurately infer chromosomal haplotypes using three or a few more single gametes and is computationally much more efficient than *Hapi*. Given that crossovers are very rare, three gametes are theoretically sufficient for phasing the entire genome, likely pushing the boundary to its possible limit. However, phasing errors may arise if two gametes have a common crossover at the same locus, which is an even rarer occurrence in practice. To deal with this possible but unlikely challenging situation, we suggest using 5 or 6 gametes rather than 3 and propose a *k*-gamete strategy, where *k* is the number of gametes and *k* > 3. The same 3-gamete core algorithm is repeatedly applied to each combination set of 3 gametes from *k* gametes, resulting in multiple candidate chromosomal haplotypes. Consensus haplotypes are then derived with high-level of confidence, ruling out the potential adverse influences that stem from (1) a common crossover shared by two gametes and/or (2) occasionally mistyped hetSNPs. In the near future, these two defect sources will likely be eliminated from the rapidly advancing technologies by further increasing the genetic marker density (DNA resolution) and decreasing the genotyping error rate, further perfecting the performance of *IIIandMe*. We foresee that *IIIandMe* will find its way to become impactful in many genetic research areas and applications.

### Data Availability

*IIIandMe* is an R package that is freely available at https://github.com/Jialab-UCR/IIIandMe.
